# Mechanical unloading coupled with coronary reperfusion reduces fibrosis and stimulates cardiomyocyte proliferation after myocardial infarction

**DOI:** 10.1101/2025.04.25.650549

**Authors:** Sean O. Bello, Charanjit Singh, Filippo Perbellini, Prakash P. Punjabi, Cesare M. Terracciano

**Affiliations:** National Heart and Lung Institute; Imperial College London - Hammersmith Campus; Imperial College London

**Author notes:** Corresponding author: Professor Cesare Terracciano. Address: Imperial Centre for Translational and Experimental Medicine, Hammersmith Hospital, National Heart & Lung Institute, Imperial College London, W12 0NN.

**Keywords:** Unloading, infarction, regeneration, fibrosis, reperfusion

## Abstract

**Introduction:** Percutaneous left ventricular assist devices (pLVADs) have become essential tools during coronary reperfusion in high risk PCI. Significant reduction in infarct propagation is observed when mechanical unloading is coupled with reperfusion, but little is known of the effect this reduction in wall stress and extracellular matrix with pLVADs have on the hearts regenerative capacity.

**Objective:** This study investigates the effect coronary reperfusion coupled with mechanical unloading has on myocardial fibrosis, and the impact these changes in extracellular matrix have on the hearts regenerative potential.

**Methods:** MI was induced by coronary artery ligation in Lewis rats. Hearts underwent permanent coronary ligation (AMI) or were reperfused after 90 minutes (AMI/R). In each group, hearts were either loaded (AMI-L or AMI/R-L) or unloaded (AMI-U or AMI/R-U). In the unloaded subgroup, the infarcted hearts were explanted after 90 minutes and transplanted into the abdomen of healthy recipients via heterotopic abdominal heart-lung transplantation. The recipient’s heart acted as control. Hearts were analysed on day 7.

**Results:** 30 hearts were studied. In the permanent ligation group, fibrosis increased in both the loaded and unloaded hearts with no significant rise in cardiomyocyte proliferation. After coronary reperfusion, there was a decrease in fibrosis with mechanical unloading and cardiomyocyte proliferation rose significantly (AMI/R-L vs AMI/R-U p= 0.0001).

Cardiomyocyte proliferative rate in the loaded and unloaded hearts was 0.6%, and 3.7% respectively after permanent ligation, and 0.5%, and 10.4% respectively after coronary reperfusion.

**Conclusion:** These data show that coronary reperfusion coupled with mechanical unloading reduces myocardial fibrosis and upregulates cardiomyocyte proliferation after myocardial infarction.

**TRANSLATIONAL PERSPECTIVE:** This study demonstrates that the adult heart’s intrinsic regenerative capacity is significantly augmented when coronary reperfusion is coupled with mechanical unloading after MI in an animal model. This finding has significant translational implications for myocardial self-repair and recovery especially in ischaemic cardiomyopathy. The investigation of cardiomyocyte proliferative response should be included in current ongoing trials such as the PROTECT IV, RECOVER IV, and IMPACT trials, all currently investigating clinical outcomes when mechanical unloading is coupled with coronary reperfusion after MI.

## INTRODUCTION

Since the second half of the 20th century, remarkable progress has been made in the treatment of cardiovascular diseases, in particular, coronary artery disease, with several major breakthroughs in the treatment for acute myocardial infarction (AMI), such as the introduction of the coronary care unit, pharmacological reperfusion (i.e., thrombolysis), pharmacological interventions (e.g., beta blockade, ACE inhibitors, antiplatelet drugs, and statins), and improvements in interventional cardiology i.e., primary percutaneous coronary intervention (PCI). ^1^ ^2^

As a result, our ability to provide patients with a viable less invasive alternative to surgical revascularisation (the gold standard for definitive treatment of coronary artery disease) has developed significantly. However, this achievement has come at the expense of a vast increase in patients with congestive heart failure (CHF), a chronic consequence of AMI. Despite the improvements in the management of acute ST elevation MI with rapid coronary reperfusion via balloon angioplasty and stenting, achieving a door-to-balloon time of <90 minutes to limit myocardial injury, nearly 10% of AMI subjects die during their index hospitalization, and about 25% of survivors progress to develop ischaemic cardiomyopathy, the leading cause of heart failure in developed countries. ^3^ ^4^

Thus, there remains a need to further optimise the management of AMI. The novel approach of coupling percutaneous left ventricular assist devices (LVADs), such as the Impella 2.5 and Impella CP, with coronary reperfusion during high risk PCI has shown improved outcomes, with studies reporting significant reduction in infarct size with unloading. ^5^ ^6^ ^7^ ^8^

This reduction in the propagation of post-infarct myocardial injury when reperfusion is coupled with mechanical unloading has been attributed to a reduction in myocardial oxygen consumption and the amelioration of ischaemia-reperfusion injury with LVAD use. ^9^

LVADs were originally developed to provide circulatory support to critically ill heart failure patients as a bridge to cardiac transplantation (BTT), bringing about significant improvements in end-organ function, haemodynamics, overall functional status and quality of life. ^10^ ^11^ Their application has however progressed beyond BTT to Destination Therapy (DT), and even Bridge To Recovery (BTR). ^12^

Temporary LVADs (Impella) are intravascular microaxial blood pumps that are miniaturised iterations of durable LVADs (e.g. HeartMate III), and are used during high risk PCI or for the management of cardiogenic shock following AMI or cardiac surgery. They include left sided devices such as the Impella 2.5, CP, 5.0, LD, and 5.5, that are positioned across the aortic valve into the left ventricle (LV) and mechanically unload the LV by providing continuous antegrade blood flow from the LV into the ascending aorta, thus reducing the workload of the LV and increasing cardiac output.

The potential for LVADs to facilitate myocardial recovery after myocardial injury is an area of much research interest. ^13^ ^14^ ^15^ By reducing myocardial wall stress via mechanically unloading the failing heart, improving cardiac excitation-contraction coupling, cardiac output and thus multi-organ function, and ameliorating the toxic neurohumoral milieu around the heart, LVADs are capable of stimulating significant improvement in myocardial function, a phenomenon described as reverse remodelling. ^16^ ^17^

Reverse remodelling refers to the termination or reversal of the pathophysiological processes that occurs following myocardial injury; it is associated with changes in the myocardium at multiple levels: whole organ, tissue, cellular, and molecular. The “main players” in the process of reverse remodelling include the cardiomyocytes, the extracellular matrix (ECM), and the microvascular system, as well as neuroendocrine mechanisms. The impact of cardiomyocyte regeneration in reverse remodelling is yet to be explored but might play a key role in the myocardial recovery observed with LVADs. ^18^

A well held paradigm up until the early parts of this century was that the heart was a post-mitotic organ and all cardiac myocytes were terminally differentiated and thus incapable of re-entering the cell cycle. In addition, most believed that there were no stem and/or progenitor cells in the heart that can differentiate into functional cardiac myocytes. More recently however, a number of remarkable studies have provided evidence to the contrary. ^19^ ^20^ Various research groups now show that the adult mammalian heart, by itself, possesses an intrinsic form of cellular homeostasis that permits regeneration and formation of new cardiac myocytes and vasculature. ^2^ ^21^ ^22^ ^23^ ^24^

Furthermore, studies in several animal models have shown that cardiomyocytes at the periphery of MI undergo endomitosis and polyploidization instead of conventional mitosis ^25^ with a majority of these cardiomyocytes harbouring a polyploid as opposed to the normal diploid genome. Interestingly, upon relief of wall stress in these hearts via mechanical unloading, there was a more than 2 fold increase in diploid cardiomyocytes with a significant decrease in the number of polyploid cardiomyocytes. It appears that the vast polyploidy of cardiomyocytes in the injured heart is a result of hypertrophic stimuli being associated with repeated rounds of DNA synthesis with an incomplete attempt to enter mitosis that has been blocked after completion of DNA replication (endomitosis). When hemodynamic stress is relieved by LVAD and the hypertrophic stimulation is decreased, this blockade is also released, resulting in karyokinesis and the formation of binucleated cardiomyocytes, cytokinesis, and real cell duplication. ^25^

Hence the favourable conditions during LVAD support, which reduces wall stress and improves the neurohormonal environment, promotes conditions for cardiac regeneration. ^26^ To date however, the impact hemodynamic unloading after MI has on the heart’s regenerative capacity remains unclear. Thus the effect of unloading coupled with coronary reperfusion after MI on load-sensitive cell-intrinsic signalling pathways such as the Hippo pathway via the expression of phosphorylated YAP (pYAP) will be described. ^27^ ^28^ ^29^

Finally, a number of contemporary studies have identified that the cardiac extracellular matrix (ECM), and its role in the propagation of myocardial fibrosis has a significant influence on the degree of reverse remodelling that occurs during mechanical unloading with LVADs. ^30^ ^31^ There are reports of fibrosis increasing during mechanical unloading in some settings which is thought to be related to the effects of unloading in reducing myocardial oxygen consumption (MV0_2_) after MI with consequent reduction in the autoregulatory stimulus necessary to generate increase in microvascular blood flow. ^32^ ^33^ Here we hypothesise that by coupling mechanical unloading with return of blood flow through the ligated major coronary vessel, these effects on microvasculature and fibrosis will be less significant. To determine ECM propagation when unloading is coupled with coronary reperfusion, the expression profile of alpha smooth muscle actin (αSMA), is investigated.^34^ ^35^

In this study, using rodent models of AMI, coronary reperfusion, and mechanical unloading via heterotopic abdominal heart and lung transplantation (HAHLT), we investigated the effect of mechanical unloading, coupled with coronary reperfusion, on the progression of myocardial fibrosis and the cardiomyocyte proliferative capacity of the infarcted heart.

## MATERIALS and METHODS

### AMI, Coronary reperfusion and Heterotopic abdominal heart & lung transplantation models

Personal, project, and establishment animal licences were granted by the UK home office as required by the Animals Scientific Procedures Act (ASPA) 1986. All animal procedures were performed in accordance with the National Institute for Health (NIH) Guide for the Care and Use of Laboratory Animals (National Research Council 2011). In brief, syngeneic male Lewis rats were anaesthetised using the inhalation anaesthetic agent isoflurane at 3-5% for induction & 1.5-2.5% for maintenance. The anaesthetic circuit used was the Bain’s co-axial facemask. Oxygen flow was kept at 3L/min for induction and 1.5-2L/min for maintenance.

Animals were then intubated using a 16 G vascular cannula and general anaesthesia achieved via a volume controlled rodent ventilator (Model 683 Rodent ventilator, Harvard 101Apparatus Ltd, UK). The tidal volume was kept at about 6-8ml/kg with a ventilatory rate of about 40 breaths/min and body temperature of 37°C. Intraoperative pain relief consisted of a single subcutaneous dose of vetergesic (buprenorphine) at 0.12mg/kg. Adequate hydration was achieved by giving 1ml bolus of warm saline subcutaneously every hour.

#### Permanent coronary artery ligation & ischaemia-reperfusion

Once adequacy of anaesthesia was confirmed by loss of pedal reflexes, the animal’s chest wall was shaved, prepped and draped, a left anterolateral thoracotomy was made over the fourth rib, and the proximal left anterior descending artery (LAD) was ligated using 6/0 prolene sutures.

After coronary artery ligation, visual assessment of the apex of the heart was carried out to determine onset of the infarct. The lungs were inflated and the chest wall temporarily closed. Animals were then left on the ventilator for 90 mins. This was taken as sufficient time to allow transmural myocardial infarction to occur.

In the loaded permanently ligated subgroup (AMI), the animal was recovered after 90 mins. In the loaded reperfusion subgroup (AMI/R-L), the chest wall was reopened after 90 mins and the ligature to the proximal LAD released to return blood flow to the infarct territory.

The chest wall was then closed and the animal recovered.

In the unloading subgroup (AMI/U or AMI/R-U), after 90 mins the torso was prepped with povidone iodine and draped. The abdominal aorta was then accessed via a midline laparotomy and adequate heparinisation was achieved by injecting heparin at 10000 IU/Kg into the inferior vena cava (IVC) using a 30G needle. The donor abdominal aorta was cannulated with a 20-gauge arterial line (Leadercath arterial, Vygon UK Ltd, FSQ049) using a catheter-over-wire technique and this was used to inject 50mls of cold St Thomas II cardioplegia solution into the donor circulation thus arresting the donor heart (permanently ligated or AMI/R) in diastole. The IVC was simultaneously transected to prevent overload and euthanasia was achieved via exanguination. The sternum was cut open and the heart explanted after ligating the IVC and superior vena cava (SVC) using 4/0 Mersilk sutures. The heart was then implanted into the abdomen of a syngeneic recipient via HAHLT.

#### Mechanical unloading (HAHLT)

The heterotopic abdominal heart lung transplantation technique was used as the model of mechanical unloading.

HAHLT in rats is a well-established technique that had been initially employed to facilitate studies in cardiac transplant immunology but has also been optimised to study the effects of mechanical unloading on the failing heart. ^36^ ^37^ ^38^ ^39^ It involves anastomosing the heart and lung en bloc from a donor onto the abdominal aorta of a healthy recipient. This model allows a much less profound or partial unloading of the LV to be achieved which is a more physiological representation of LVAD-induced mechanical unloading.^40^ Only a single anastomosis of the ascending aorta of the donor heart to the abdominal aorta of the recipient is completed. As the donor IVC and SVC are ligated and the pulmonary circulation remains intact, the LV receives additional blood from the right ventricle (RV), via the pulmonary veins after passing through the lungs. (Illustrated in figure S1, supplementary page 4).

The surgical steps have been previously described. In brief, both the recipients’ and donor torsos were shaved, prepped with povidone iodine and draped as shown in Figure 1 below. The recipients’ abdominal aorta was accessed via a midline laparotomy incision and with the aid of a microscope, a 0.5cm midline longitudinal aortotomy was carried out and prepared for anastomosis after achieving proximal and distal vascular control. The donor abdominal aorta was also accessed and the donor heart explanted as described above.

**Figure 1:**
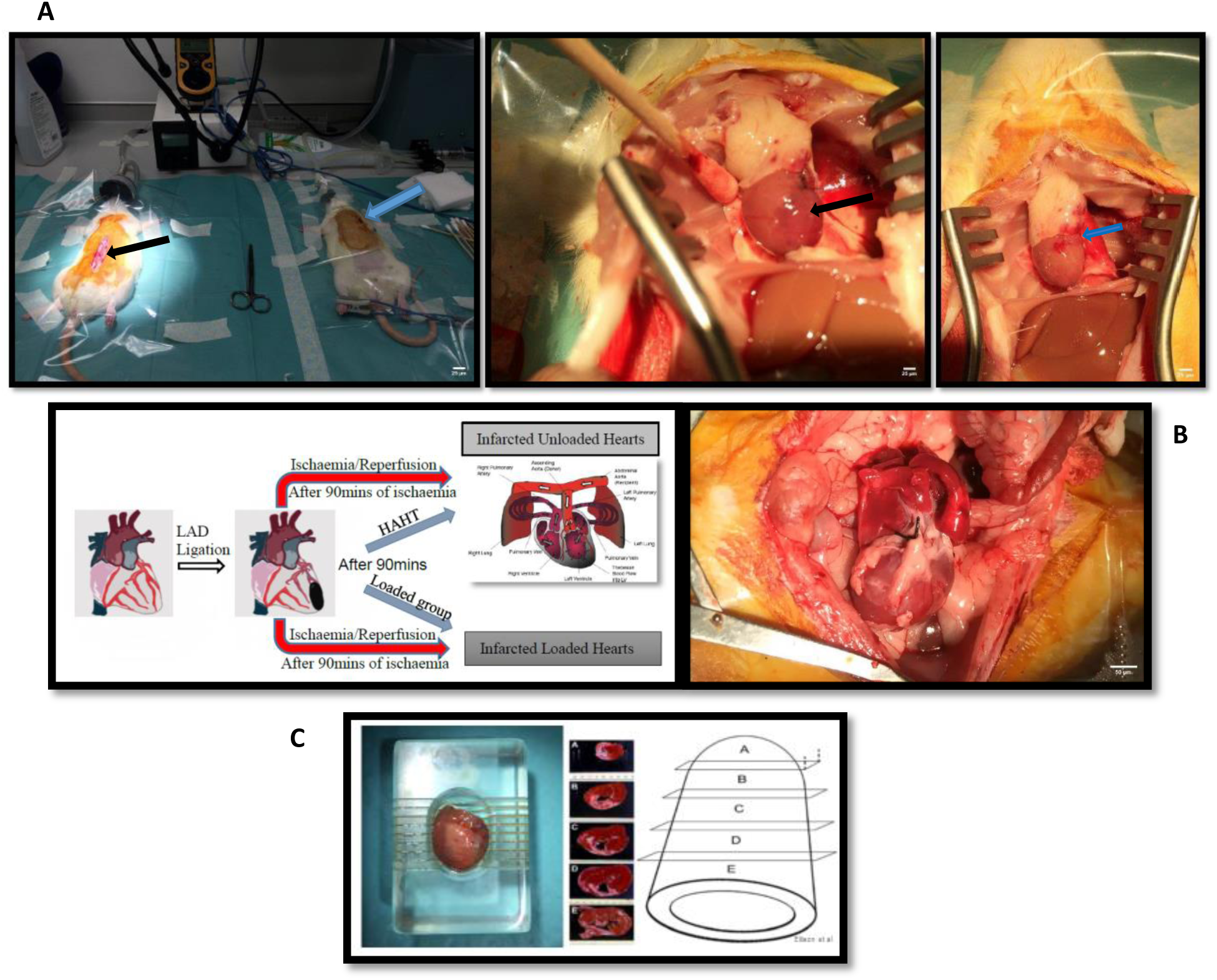
Representative images summarising the procedural steps involved in animal model creation. (A) After 90 minutes of MI, hearts were either left with permanent coronary artery ligation or reperfused by ligature release. Rats were then either recovered (loaded group) or hearts were explanted and implanted into the abdomen of a healthy recipient for mechanical unloading via HAHLT. Note the blue arrow in the left image in (A) pointing at the closed left thoracotomy wound following coronary ligature. The rat on the left is being prepared for HAHLT with the black arrow pointing at the laparotomy incision. The middle image in (A) shows the ligated heart after undergoing diastolic arrest following infusion of cold crystalloid cardioplegia solution prior to explantation for HAHLT, note the black arrow pointing at the unblanched infarcted territory downstream from the LAD ligature (blue arrow). The right image in (A) shows the previously well demarcated infarct territory has now been blanched (blue arrow) following ligature release and infusion of crystalloid cardioplegia, indicating adequate coronary reperfusion. Left image in (B) is a schematic representation of the steps described in (A) and includes a graphic illustrating HAHLT. In HAHLT, the ascending aorta of the donor heart is anastomosed to the abdominal aorta of the recipient and as the SVC and IVC are ligated, only the coronary flow down the donor’s aorta returns to the LV. The right image in (B) shows the donor heart in the abdomen of the recipient, note the blue arrow pointing at the IVC ligature. Following heart harvest, the atria were excised and the ventricles placed in an acrylic zivic rat heart slicer matrix (C) where five 2mm coronal section slice intervals (A-E) were obtained for analysis. Right sided image of (C) reproduced with permission from Ellison et al 2011. ^24^

#### Animal recovery and post-operative care

Recovered animals were observed in the operating theatre for an hour following weaning from general anaesthesia at which point signs of pain or distress were acted upon in accordance with Home Office regulations. Post-operative analgesia was achieved by injecting carprofen (rimadyl) subcutaneously at 0.5mg/kg at the end of the operation and once daily for two days. Antibiotic cover consisted of enrofloxacin given subcutaneously at 1mg/kg at the end of the operation and daily for two days. Animals were checked twice a day for the first 3 post-operative days. Daily weights and health checks were performed thereafter.

30 male syngeneic Lewis rats weighing 250-300g (Charles River, UK) were utilised to create models of AMI, coronary reperfusion (R), and mechanical unloading (U) via HAHLT.

Detailed description of the operative procedures including heart harvest on day 7, tissue preparation and experimental techniques are provided in the supplementary pages (2-16).

### Animal models & experimental groups

As described above, MI was induced by either permanently ligating the LAD (AMI) or reperfusing the vessel after 90 minutes (AMI/R). In each group hearts were either loaded (AMI-L or AMI/R-L) or unloaded (AMI-U or AMI/R-U). The animals of the loaded subgroup were recovered, whilst in the unloaded subgroup, the infarcted hearts were explanted after 90 minutes and transplanted into the abdomen of healthy recipients via HAHLT. The recipient’s heart acted as control.

Hearts were explanted on day 7 for histology and immunohistochemistry. (Fig. 1)

4 subgroups were formed: AMI loaded (AMI-L), AMI reperfusion loaded (AMI/R-L), AMI unloaded (AMI-U), AMI reperfusion unloaded (AMI/R-U).

Detailed annotations for each group are provided in the supplementary pages (9-10). All animals were randomly assigned identification codes and these were blinded to the primary investigator (first author) who carried out all the experiments as detailed in the supplementary pages (10-16).

### Histological analysis of frozen tissue sections for the quantification of myocardial fibrosis

15μm cryosections were prepared and processed with histological stains to assess the degree of fibrosis in the rat models of MI. Sirius red/Fast green collagen staining was employed using a standardised protocol. Images of the stained sections were obtained with widefield microscopy and with the aid of imageJ software, infarct size (IS) and area at risk (AAR) were determined using planimetry. (Illustrated in figure S6A & S6B, supplementary page 12)

### Mode of detection of proliferating cells

Immunohistochemistry of frozen cross sections using the proliferative marker Ki67 was employed to determine cell proliferation. Ki67’s sole expression during the active phases of the cell cycle (G1, S, G2, and mitosis) distinguishes it from markers that are heavily influenced by DNA damage and repair such as the proliferating cell nuclear antigen or the thymidine analogue Bromodeoxyuridine (BrdU). ^41^ ^42^

### Western Blotting

Western blotting technique was used to determine the expression profiles of αSMA and pYAP.

Details of the above methods are provided in the supplementary pages (2-15).

### Statistical analysis

The degree of myocardial fibrosis and cardiomyocyte proliferation was compared between the loaded and unloaded permanently ligated- and ischaemia-reperfused- hearts. A two-tailed t-test was used for the comparison between independent groups of normally distributed data as determined by Kolmogorov-Smirnov testing. One-way analysis of variance (ANOVA) followed by Bonferroni post hoc test for individual significant differences was used. All statistical analyses were done using Prism 8 software (GraphPad Software, Inc.). Five animals (n=5) were studied in each subgroup. The results are presented as means ±SD. The significance threshold was set at p<0.05 for all data. * denotes P < 0.05, ** denotes P < 0.01 and *** denotes P < 0.001.

## RESULTS

### Effect of Mechanical Unloading on Myocardial Fibrosis after Acute Myocardial Infarction

After AMI, mechanical unloading in the permanently ligated hearts was associated with a statistically significant increase in fibrosis compared to loaded hearts at day 7 (AMI-L vs AMI-U p= 0.0363). However, in the coronary reperfusion group, mechanical unloading after acute MI was not associated with increase in fibrosis (AMI/R-L vs AMI/R-U p= 0.5711) as illustrated in figures 2 & 3.

**Figure 2:**
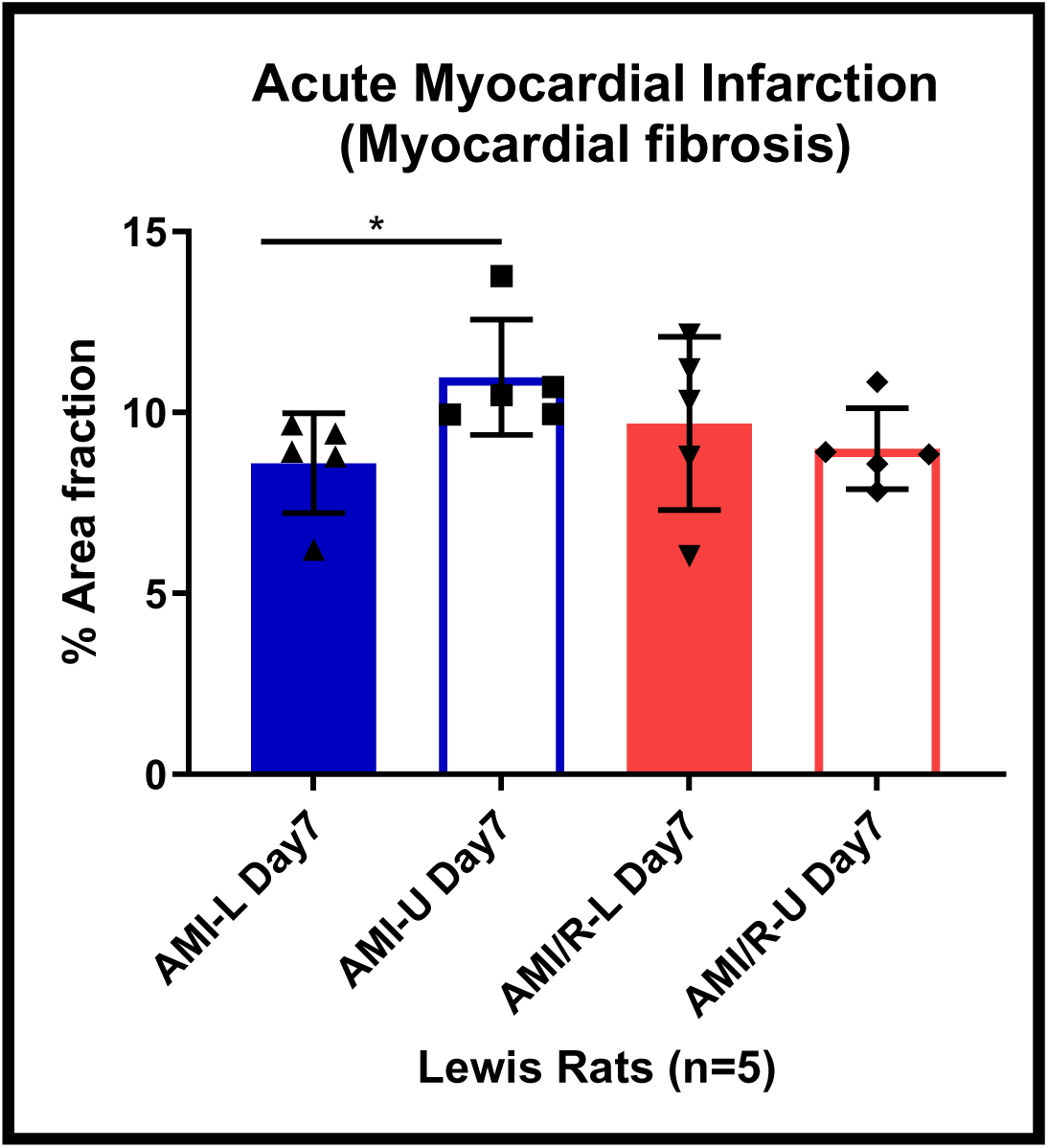
Differences in myocardial fibrosis between loaded and unloaded hearts after permanent coronary artery ligation and coronary reperfusion. Note the significant increase in fibrosis noted after mechanical unloading in the permanently ligated hearts was not observed in the mechanically unloaded reperfused hearts. Blue bars represent permanently ligated hearts while red bars are the reperfused hearts. The full bars are the loaded hearts while the empty bars represent the mechanically unloaded hearts. There were 5 rat hearts studied per group. One-way analysis of variance (ANOVA) followed by Bonferroni post hoc test for individual significant differences was used. All statistical analyses were done using Prism 8 software (GraphPad Software, Inc.). The results are presented as means ±SD. The significance threshold was set at p<0.05 for all data. * denotes P < 0.05, ** denotes P < 0.01 and *** denotes P < 0.001.

In the unloaded reperfused hearts, areas furthest from the infarct site showed less fibrosis (Fig. 3D) compared to that observed in the unloaded permanently ligated hearts (Fig. 3B).

**Figure 3:**
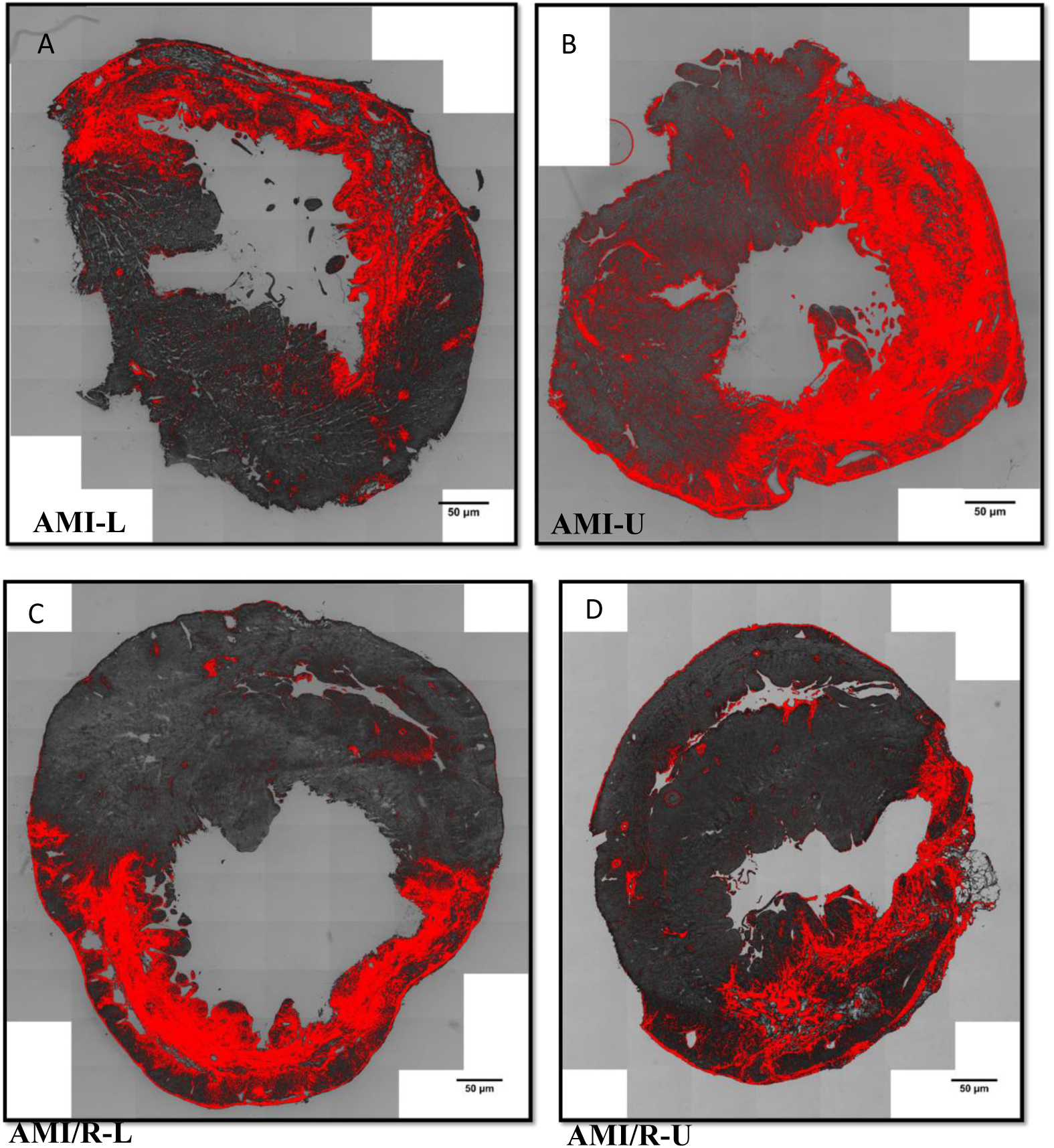
Histological samples at day 7. (A) Myocardial fibrosis in AMI-L with increase in fibrosis evident after mechanical unloading in the AMI-U (B). In the coronary reperfusion group (C-D) however, mechanical unloading in the AMI/R-U (D) did not significantly change the extent of fibrosis compared to that in the loaded AMI/R-L heart (C). Areas of fibrosis was determined by Sirius red staining and quantified using ImageJ software.

### Regional Evaluation of Cardiomyocyte Proliferation in Permanently Ligated and Coronary Reperfused Hearts

Analysis of cardiomyocyte proliferation was completed at day 7 as several studies have reported significant cardiomyocyte regenerative responses to myocardial injury within this time point.^43^ ^44^ ^45^ Regional analysis of proliferation was completed to facilitate the determination of the differential effects of mechanical unloading on cardiomyocyte proliferation after MI in areas of the myocardium close to the primary infarct compared to more distant myocardium. The regions assessed are illustrated in figure 4A.

**Figure 4:**
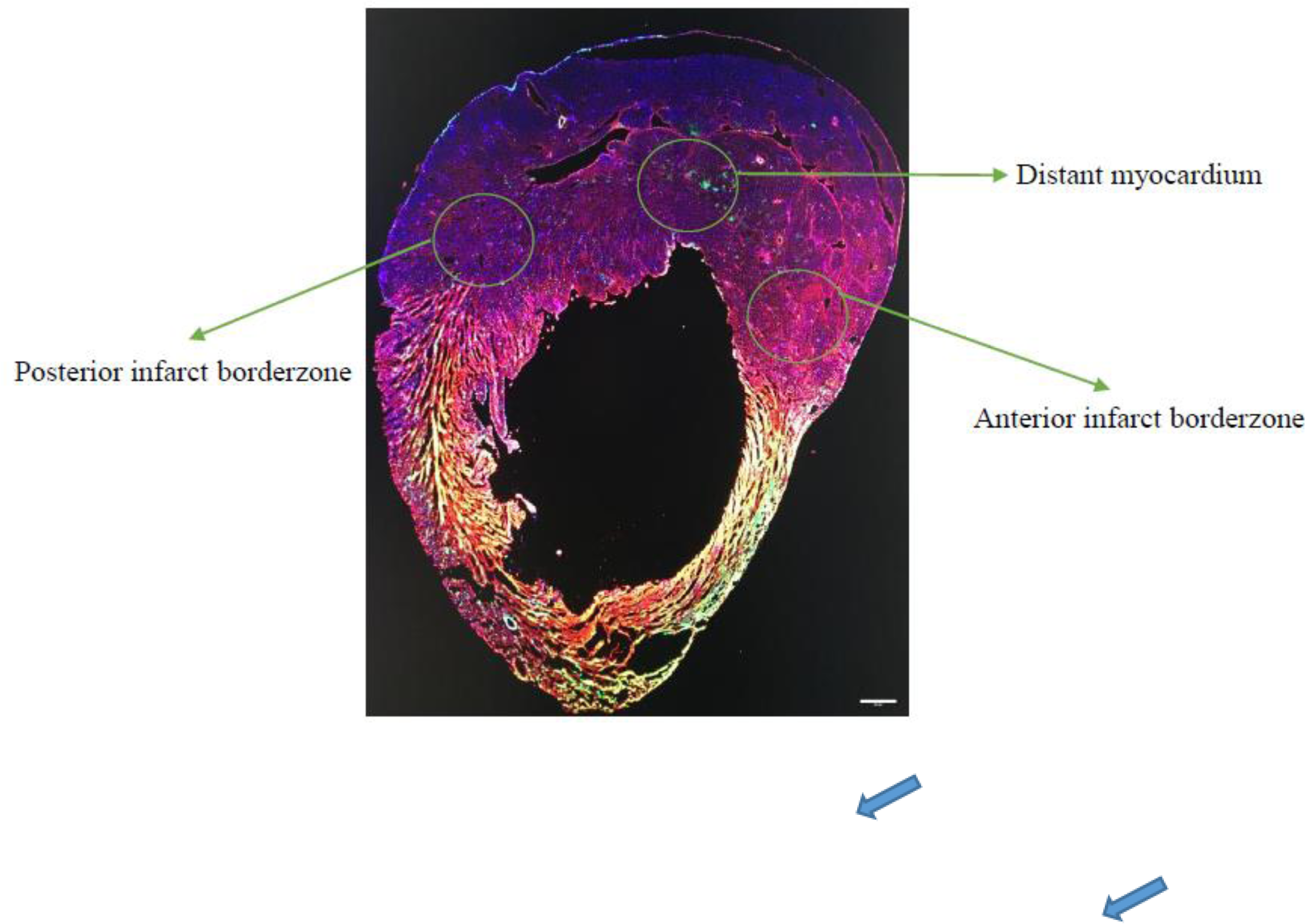
Bright field microscopy (A) of a 15μm frozen cross section of a Lewis rat heart after immunohistochemistry staining using cTnT (red) to identify cardiomyocytes, DAPI (blue) for the nuclei, and Ki67 (green) to detect proliferating cells. The three regions evaluated to determine cardiomyocyte proliferative response to MI are shown. (B) Confocal microscopic image of the tissue cross section showing 2 proliferating cardiomyocytes (blue arrows).

The anterior infarct borderzone represents the area between the anterior wall of the right ventricle, septum, and the anterior aspect of the LV free wall. The posterior infarct borderzone is the area between the posterior wall of the right ventricle, septum, and the posterior aspect of the left ventricular free wall. Finally, the distant myocardium is the midpoint of the interventricular septum. Cardiac troponin (cTnT) positive cells (stained red in Fig. 4B) expressing Ki67 (green) in their nuclei (blue) were counted as proliferating cardiomyocytes.

#### Anterior infarct borderzone

In the anterior infarct borderzone at day 7, cardiomyocyte proliferation was 4 times higher in the loaded infarcted hearts compared to control after permanent coronary artery ligation (AMI-L vs CTR p=0.0185). After mechanical unloading, there was no statistically significant difference in cardiomyocyte proliferation between the loaded and unloaded hearts in the anterior infarct borderzone after permanent coronary artery ligation. (Fig. 5A)

**Figure 5:**
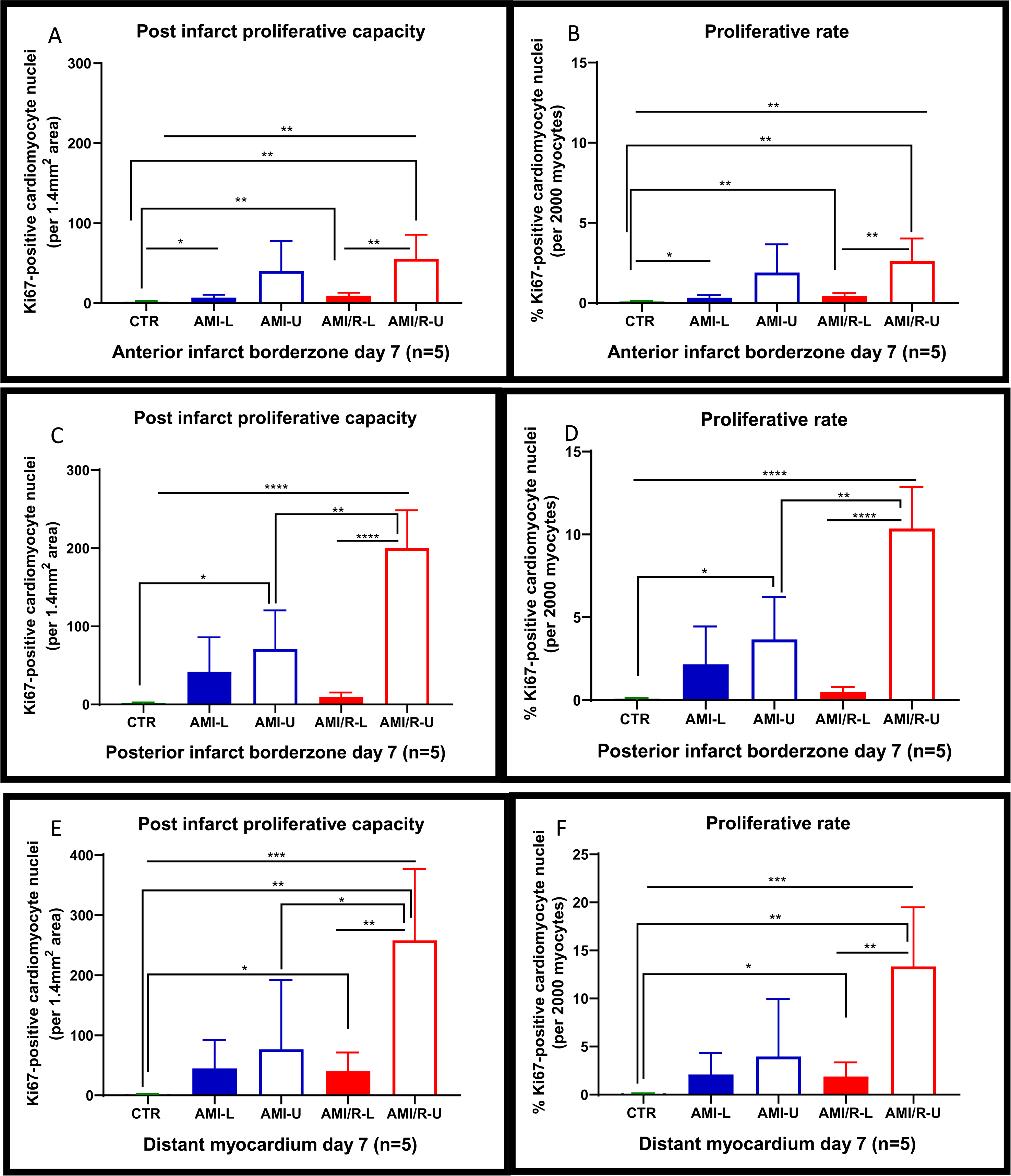
(A-F) Regional proliferation of Ki67-positive cardiomyocytes in the anterior and posterior infarct borderzone, and distant myocardium 7 days after permanent coronary artery ligation or coronary reperfusion. In each illustrative figure, the filled bars represent loaded hearts and the empty bars unloaded hearts. The permanently ligated coronary artery group are in blue and the coronary reperfusion group in red. The proliferative capacity reflects the total number of Ki67-positive cardiomyocyte in the chosen region of myocardium. The proliferative rate is the number of proliferating cardiomyocytes per 2000 cardiomyocytes in each chosen region. One-way analysis of variance (ANOVA) followed by Bonferroni post hoc test for individual significant differences was used. All statistical analyses were done using Prism 8 software (GraphPad Software, Inc.). The results are presented as means ±SD. The significance threshold was set at p<0.05 for all data. * denotes P < 0.05, ** denotes P < 0.01 and *** denotes P < 0.001.

After coronary reperfusion, there was a statistically significant 5 times increase in cardiomyocyte proliferation in the loaded hearts compared with control (AMI/R-L vs CTR p= 0.0027). Mechanical unloading coupled with coronary reperfusion stimulated a significant increase in cardiomyocyte proliferation compared to both the control hearts (AMI/R-U vs CTR p= 0.004) and the loaded reperfused hearts (AMI/R-U vs AMI/R-L p=0.0092) as shown in figure 5A. The proliferative rate in the control, loaded, and unloaded hearts was 0.1%, 0.3%, and 2% in the permanently ligated hearts, and 0.1%, 0.4%, and 2.6% in the reperfused hearts respectively. (Fig. 5B)

#### Posterior infarct borderzone

In the posterior infarct borderzone at day 7, permanent coronary artery ligation was associated with an increase in cardiomyocyte proliferation in the loaded hearts compared to control but this was not statistically significant (AMI-L vs CTR p=0.0781). (Fig. 5C)

There was also no statistically significant difference in cardiomyocyte proliferation between the loaded and unloaded permanently ligated hearts. (Fig. 5C) In the coronary reperfusion group however, there was a statistically significant 5 times increase in cardiomyocyte proliferation in the loaded hearts compared to control (AMI/R-L vs CTR p=0.0118). After mechanical unloading coupled with coronary reperfusion, cardiomyocyte proliferation rose quite significantly compared to the loaded reperfused hearts (AMI/R-U vs AMI/R-L p= 0.0001). (Fig. 5C). Mechanical unloading coupled with coronary reperfusion was also associated with a significant increase in proliferation when compared to unloading alone without reperfusion (AMI/R-U vs AMI-U p= 0.0031). The proliferative rate in the control, loaded, and unloaded hearts was 0.1%, 0.6%, and 3.7% in the permanently ligated hearts, and 0.1%, 0.5%, and 10.4% in the reperfused hearts respectively. (Fig. 5D)

#### Distant myocardium

In the distant myocardium at day 7, there was also no statistically significant difference in cardiomyocyte proliferation between the loaded and unloaded hearts after permanent coronary artery ligation. (Fig. 5E) After coronary reperfusion however, there was a statistically significant increase in cardiomyocyte proliferation in the loaded hearts compared

to control (AMI/R-L vs CTR p= 0.0248) and this increase was even larger after mechanical unloading (AMI/R-U vs CTR p= 0.0013). Mechanical unloading coupled with coronary reperfusion resulted in a large and statistically significant increase in cardiomyocyte proliferation compared to the loaded reperfused hearts (AMI/R-U vs AMI/R-L p= 0.0042). In addition, there was a statistically significant increase in proliferation in the unloaded reperfused hearts compared to the unloaded permanently ligated hearts (AMI/R-U vs AMI-U p= 0.0402). (Fig. 5E)

The proliferative rate in the control, loaded, and unloaded hearts was 0.1%, 2.1%, and 3.9% in the permanently ligated hearts, and 0.1%, 1.9%, and 13.4% in the reperfused hearts respectively. (Fig. 5F)

### Effect of Mechanical Unloading on the Dynamic Expression Profiles of αSMA and pYAP after Myocardial Infarction

#### Effect of mechanical unloading on αSMA expression after acute myocardial infarction

In the loaded permanently ligated heart, AMI resulted in a statistically significant rise in αSMA protein expression compared to control at day 7 (AMI-L vs CTR p= 0.0291) as shown in figure 6A below. Mechanical unloading was associated with a further rise in αSMA protein expression but this was not statistically significant. In the coronary reperfusion group, αSMA protein expression also rose significantly in the loaded hearts compared to control (AMI/R-L vs CTR p= 0.0020). However, in contrast to the permanent ligation group, mechanical unloading coupled with coronary reperfusion led to a statistically significant decrease in αSMA protein expression (AMI/R-L vs AMI/R-U p= 0.0483). (Fig. 6A)

**Figure 6:**
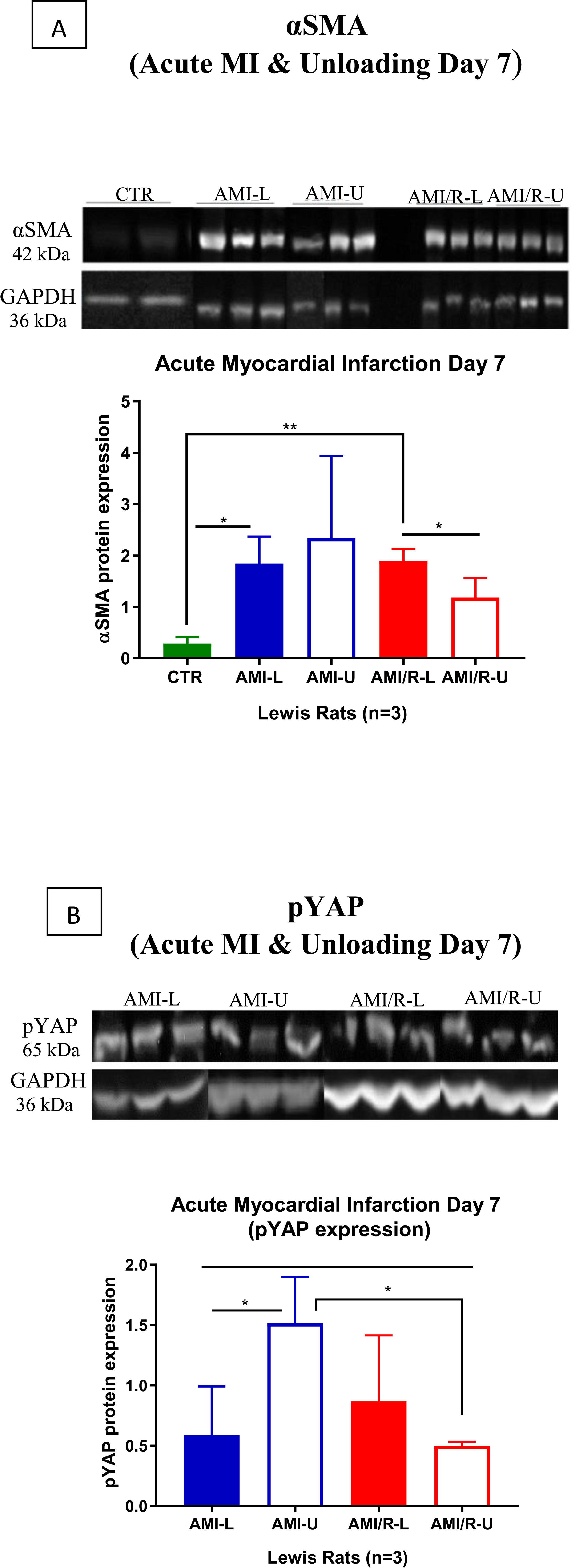
(A & B) showing the effects of mechanical unloading on the expression profiles of αSMA and pYAP after 7 days of permanent coronary artery ligation and coronary reperfusion. Three rats were tested in each group. In each illustrative figure, the filled bars represent loaded hearts and the empty bars unloaded hearts. The permanently ligated coronary artery group are in blue and the coronary reperfusion group in red. One-way analysis of variance (ANOVA) followed by Bonferroni post hoc test for individual significant differences was used. All statistical analyses were done using Prism 8 software (GraphPad Software, Inc.). The results are presented as means ±SD. The significance threshold was set at p<0.05 for all data. * denotes P < 0.05, ** denotes P < 0.01 and *** denotes P < 0.001.

**Figure 7:**
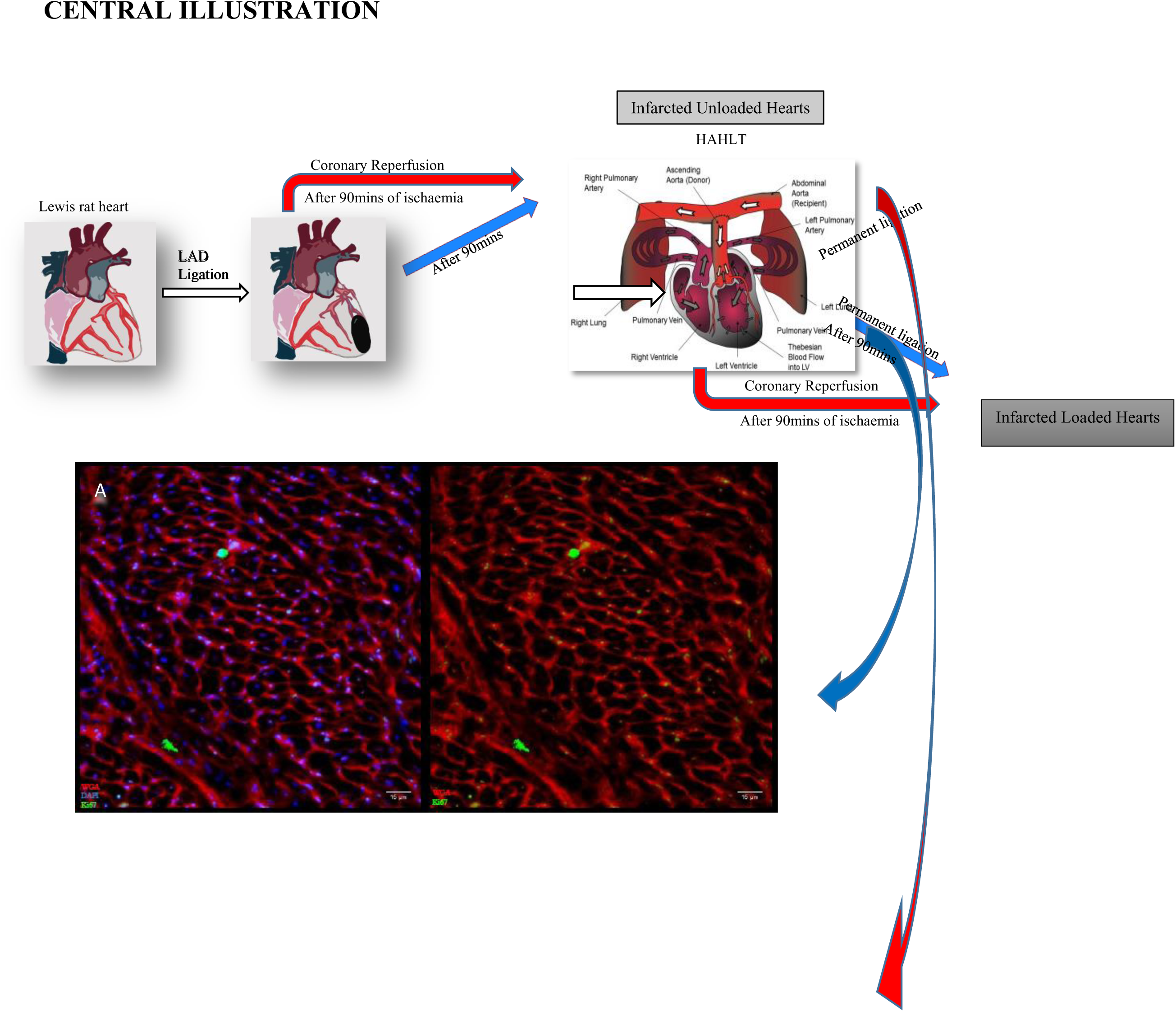
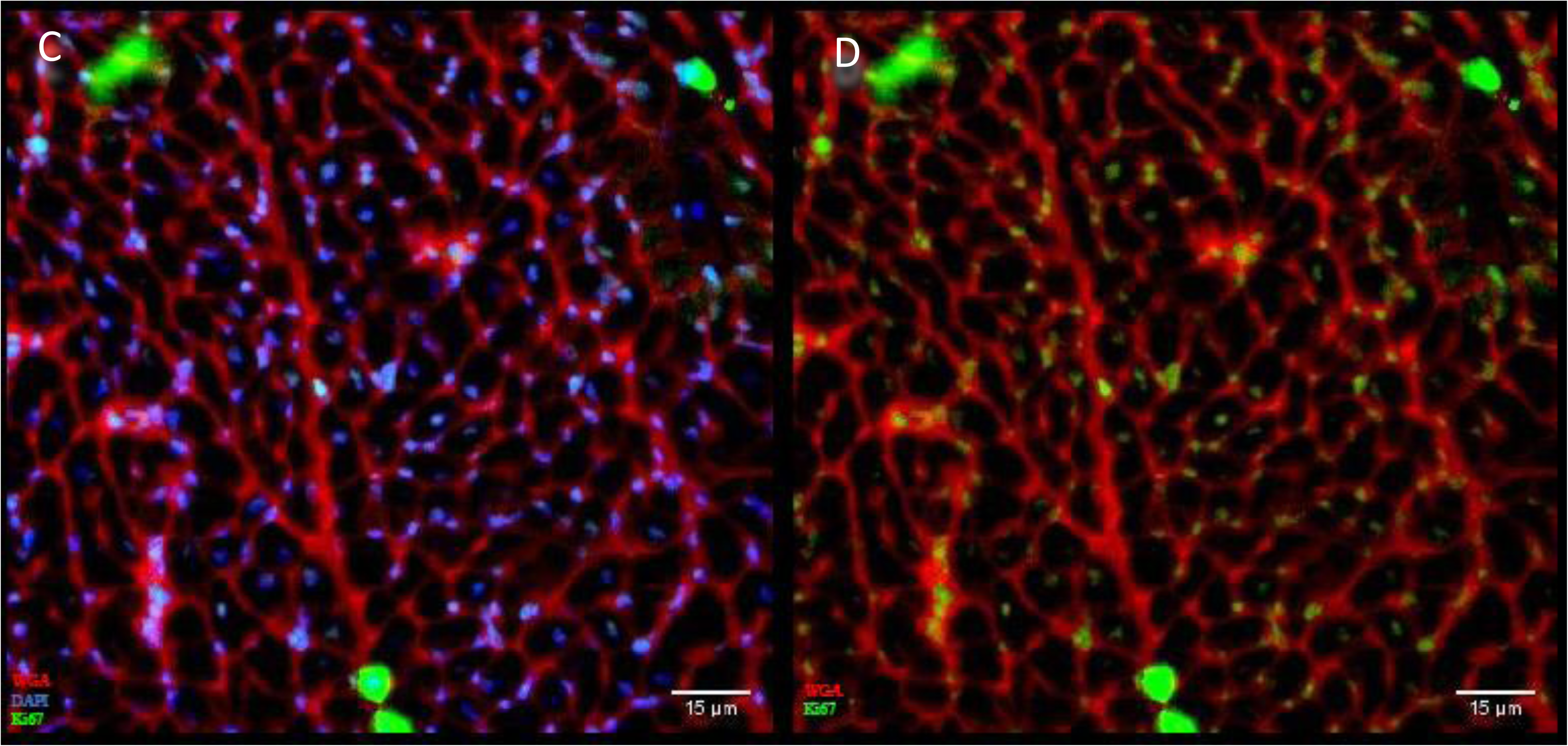
Schematic diagram showing infarcted Lewis rat hearts after either permanent LAD ligation (blue arrows) or coronary reperfusion (red arrows). Both groups were then either left loaded or the hearts explanted and mechanically unloaded via heterotopic abdominal heart and lung transplantation (HAHLT). (A-D) are immunohistochemistry stains for the quantification of cardiomyocyte proliferation. Red is the membrane marker wheat germ agglutinin (WGA). Blue is the nuclei marker (DAPI). Green is the proliferative marker (Ki67). B is the same image as A, but with the blue nuclei stain silenced to allow better visualisation of Ki67 stain. Similarly D is the same image as C but with the nuclei stain silenced. Cardiomyocyte proliferative capacity was stunted in the loaded hearts irrespective of ligated vessel reperfusion (A-B). Only when mechanical unloading was coupled with coronary reperfusion did significant augmentation of cardiomyocyte proliferation occur (C-D). Embedded HAHLT figure reproduced with permission from Ibrahim 2013. ^39^

#### Effect of mechanical unloading on pYAP expression after acute MI

At day 7 after permanent coronary artery ligation, there was a statistically significant increase in pYAP expression in the unloaded hearts compared to the loaded hearts (AMI-U vs AMI-L P=0.0450). (Fig. 6B)

However, mechanical unloading coupled with coronary reperfusion was associated with a statistically significant decrease in pYAP protein expression compared to the unloaded permanently ligated hearts (AMI/R-U vs AMI-U p=0.0104). (Fig. 6B)

## DISCUSSION

Data from this study shows that mechanical unloading coupled with coronary reperfusion is associated with an increase in the heart’s endogenous cardiomyocyte proliferative capacity. Increases in cardiomyocyte proliferative rate after myocardial injury ranged from 0.3% to 2.1% in the loaded permanently ligated and reperfused hearts. However, after mechanical unloading was coupled with coronary reperfusion, there was a much larger and statistically significant rise in proliferation of new cardiomyocytes, particularly in the regions of the myocardium furthest away from the primary area of infarct, i.e. 2.6%, 10.4%, and 13.4%, in the anterior infarct borderzone, posterior infarct borderzone, and distant myocardium, respectively.

To further assess the impact mechanical unloading coupled with coronary reperfusion has on the heart’s endogenous cardiomyocyte proliferative capacity, the effect of unloading on the activity of one of the cell-intrinsic signalling pathways that regulate the proliferation of new cardiomyocytes was investigated. The expression of YAP and pYAP impacts cardiomyocyte proliferation. ^46^ ^47^ YAP’s presence in the cell nucleus is vital for cellular proliferation. When phosphorylated, YAP becomes extruded from the nucleus and is located in the cytosol as pYAP with consequent reduction in cell proliferation. Thus high levels of pYAP are associated with a downregulation of cellular proliferation. ^46^ ^47^ ^48^ Data from this study showed that mechanical unloading coupled with coronary reperfusion resulted in a statistically significant decrease in the levels of pYAP compared to the unloaded permanently ligated hearts. This correlates well with the high levels of cardiomyocyte proliferation seen on immunohistochemistry when mechanical unloading is coupled with coronary reperfusion. In the absence of coronary reperfusion, mechanical unloading alone was associated with a statistically significant increase in the expression of pYAP corresponding with the muted regenerative response observed in this setting.

The augmented endogenous cardiomyocyte proliferative capacity observed in this study occurred in areas with decreased myocardial fibrosis. It is worth noting that myocardial fibrosis increased when the permanently ligated hearts were mechanically unloaded. There was a significant rise in fibrosis after 7 days (21%) of unloading with no significant increase in cardiomyocyte proliferation. This observed increase in fibrosis with mechanical unloading has been previously reported. ^49^ Remarkably however, when mechanical unloading was coupled with coronary reperfusion, there was no significant increase in myocardial fibrosis and cardiomyocyte proliferation rose significantly. This supports our hypothesis that coupling mechanical unloading with coronary reperfusion likely ameliorates the lost autoregulatory stimulus on the microvasculature with consequent reduction in fibrosis, and that this contributes to the generation of a neurohumoral environment conducive for cardiomyocyte proliferation.

In an attempt to better understand the acute effects of mechanical unloading on the propagation of myocardial fibrosis after permanent coronary ligation and coronary reperfusion, the expression profile of αSMA in each group was determined. At day 7, mechanical unloading coupled with coronary reperfusion led to a statistically significant decrease in αSMA levels. This suggests that the activity of myofibroblasts might be downregulated when mechanical unloading is coupled with coronary reperfusion with consequent decrease in ECM protein deposition. This might also explain the amelioration of the increase in myocardial fibrosis that is seen in the permanently ligated group after unloading.

The findings from this study supports the notion that mechanical unloading promotes the reversal of the injured adult heart back to its foetal haemodynamic phenotype with a reduction in mechanical load and myocardial wall stress and a resultant increase in its hyperplastic (as opposed to hypertrophic) growth that favours cardiomyocyte proliferation. However, in the absence of coronary reperfusion, the reduction in MV0_2_ that occurs with unloading appears to interfere with the autoregulatory stimulus that would normally augment microvascular blood flow to compensate for the loss of perfusion from an obstructed parent vessel. The reported increase in myocardial fibrosis in this setting could be hindering cardiomyocyte proliferation, myocardial repair and hence recovery. ^31^ ^32^

The synergistic effect of coronary reperfusion and mechanical unloading in reducing the propagation of myocardial fibrosis and stimulating significant proliferation of new cardiomyocytes has major clinical implications as it suggests that after acute MI, mechanical unloading at the time of primary PCI could potentiate the hearts own intrinsic capacity for self-repair. It also adds to the growing body of evidence supporting the use of short term mechanical circulatory support for the management of AMI in high risk patients undergoing PCI or surgery. Outcomes of current ongoing trials such as PROTECT IV RCT, RESTORE IV, and the IMPACT trial are much awaited. ^50^ ^51^

## STUDY LIMITATIONS

The use of Ki67 as a proliferative marker in this study is limited by the fact that whilst it identifies cells at any of the phases of the cell cycle aside from G0 (G1, S, G2 and M phases), it does not determine the presence of true cell division as opposed to binucleation or multinucleation. For this reason we only counted one nuclei per cell and only counted cardiomyocytes with well-defined borders using WGA staining. Thus we have understated the total number of possible proliferating cardiomyocytes in this study. Future studies using markers of mitosis such as Aurora B and or phosphorylated histone will be beneficial.

## CONCLUSION

The findings described in this study have significant implications for the management of acute myocardial infarction as it is the first report to show that coupling mechanical unloading with coronary reperfusion not only provides haemodynamic support for these high risk patients but also help stimulate the hearts intrinsic capacity for self-repair. Further studies are needed to corroborate these findings.

## Acknowledgement

We give thanks to Dr Christopher Kane, Dr Peter Wright, Dr Katherine Mansfield, and Peter O’Gara for their valued input and technical support throughout the study.

## REFERENCES

1. Andersson C, Nayor M, Tsao CW, Levy D, Vasan RS. Framingham Heart Study. J Am Coll Cardiol. 2021;77(21):2680–2692. doi:10.1016/j.jacc.2021.01.059

2. Koudstaal S, Jansen Of Lorkeers SJ, Gaetani R, et al. Concise review: heart regeneration and the role of cardiac stem cells. Stem Cells Transl Med. 2013;2:434–443. doi:10.5966/sctm.2013-0001

3. Kloner RA, Schwartz Longacre L. State of the science of cardioprotection: Challenges and opportunities-Proceedings of the 2010 NHLBI workshop on cardioprotection. J Cardiovasc Pharmacol Ther. 2011;16(3-4):223–232. doi:10.1177/1074248411402501

4. Pastena P, Frye JT, Ho C, Goldschmidt ME, Kalogeropoulos AP. Ischemic cardiomyopathy: epidemiology, pathophysiology, outcomes, and therapeutic options. Heart Fail Rev. 2024;29(1):287–299. doi:10.1007/s10741-023-10377-4

5. Neill WWO, Kleiman NS, Moses J, et al. A Prospective, Randomized Clinical Trial of Hemodynamic Support With Impella 2. 5 Versus Intra-Aortic Balloon Pump in Patients Undergoing High-Risk Percutaneous. Circulation. 2012;(126):1717–1727. doi:10.1161/CIRCULATIONAHA.112.098194

6. Kapur NK, Paruchuri V, Urbano-Morales JA, et al. Mechanically unloading the left ventricle before coronary reperfusion reduces left ventricular wall stress and myocardial infarct size. Circulation. 2013;128(4):328–336. doi:10.1161/CIRCULATIONAHA.112.000029

7. Navin K. Kapur M, et al. Unloading the Left Ventricle Before Reperfusion in Patients With Anterior STSegment–Elevation Myocardial Infarction. Circulation. 2019:337–346. doi:10.1161/CIRCULATIONAHA.118.038269

8. Ko B, Drakos SG, Ibrahim H, et al. Percutaneous Mechanical Unloading Simultaneously with Reperfusion Induces Increased Myocardial Salvage in Experimental Acute Myocardial Infarction. Circ Hear Fail. 2020;13(1):E005893. doi:10.1161/CIRCHEARTFAILURE.119.005893

9. Kapur NK, Qiao X, Paruchuri V, et al. Mechanical Pre-Conditioning With Acute Circulatory Support Before Reperfusion Limits Infarct Size in Acute Myocardial Infarction. JACC Hear Fail. 2015;3(11):873–882. doi:10.1016/j.jchf.2015.06.010

10. Holley CT, Harvey L, John R. Left ventricular assist devices as a bridge to cardiac transplantation. J Thorac Dis. 2014;6(8):1110–1119. doi:10.3978/j.issn.2072-1439.2014.06.46

11. Lee S, Fukamachi K, Golding L, Moazami N, Starling RC. Left Ventricular Assist Devices: From the Bench to the Clinic. Cardiology. 2013;125(1):1–12. doi:10.1159/000346865

12. Topkara VK, Garan AR, Fine B, et al. Myocardial Recovery in Patients Receiving Contemporary Left Ventricular Assist DevicesCLINICAL PERSPECTIVE. Circ Hear Fail. 2016;9(7):e003157. doi:10.1161/CIRCHEARTFAILURE.116.003157

13. Frazier OH, Benedict CR, Radovancevic B, et al. Improved left ventricular function after chronic left ventricular unloading. Ann Thorac Surg. 1996;62(3):672–675.

14. Maybaum S, Mancini D, Xydas S, et al. Cardiac improvement during mechanical circulatory support: A prospective multicenter study of the LVAD working group. Circulation. 2007;115(19):2497–2505. doi:10.1161/CIRCULATIONAHA.106.633180

15. Terracciano CMN, Hardy J, Birks EJ, Khaghani a., Banner NR, Yacoub MH. Clinical recovery from end-stage heart failure using left-ventricular assist device and pharmacological therapy correlates with increased sarcoplasmic reticulum calcium content but not with regression of cellular hypertrophy. Circulation. 2004;109:2263–2265. doi:10.1161/01.CIR.0000129233.51320.92

16. Saeed D, Feldman D, Banayosy A El, et al. The 2023 International Society for Heart and Lung Transplantation Guidelines for Mechanical Circulatory Support : A 10- Year Update. JHLT. 2023;42(7):e1-e222. doi:10.1016/j.healun.2022.12.004

17. Birks EJ, Drakos SG, Patel SR, et al. Prospective Multicenter Study of Myocardial Recovery Using Left Ventricular Assist Devices (RESTAGE-HF [Remission from Stage D Heart Failure]) Medium-Term and Primary End Point Results. Circulation. 2020;142(21):2016–2028. doi:10.1161/CIRCULATIONAHA.120.046415

18. Canseco DC, Kimura W, Garg S, et al. Human Ventricular Unloading Induces Cardiomyocyte Proliferation. J Am Coll Cardiol. 2015;65(9):1–9. doi:10.1016/j.jacc.2014.12.027

19. Hsieh PCH, Segers VFM, Davis ME, et al. Evidence from a genetic fate-mapping study that stem cells refresh adult mammalian cardiomyocytes after injury. Nat Med. 2007;13(8):970–974. doi:10.1038/nm1618

20. Bergmann O, Bhardwaj RD, Bernard S, et al. Evidence for cardiomyocyte renewal in humans. Science. 2009;324(5923):98-102. doi:10.1126/science.1164680

21. Kajstura J, Urbanek K, Perl S, et al. Cardiomyogenesis in the adult human heart. Circ Res. 2010;107(2):305–315. doi:10.1161/CIRCRESAHA.110.223024

22. Ellison GM, Nadal-Ginard B, Torella D. Optimizing cardiac repair and regeneration through activation of the endogenous cardiac stem cell compartment. J Cardiovasc Transl Res. 2012;5:667–677. doi:10.1007/s12265-012-9384-5

23. Malliaras K, Zhang Y, Seinfeld J, et al. Cardiomyocyte proliferation and progenitor cell recruitment underlie therapeutic regeneration after myocardial infarction in the adult mouse heart. EMBO Mol Med. 2013;5(2):191–209. doi:10.1002/emmm.201201737

24. Ellison GM, Torella D, Dellegrottaglie S, et al. Endogenous cardiac stem cell activation by insulin-like growth factor-1/hepatocyte growth factor intracoronary injection fosters survival and regeneration of the infarcted pig heart. J Am Coll Cardiol. 2011;58(9):977–986. doi:10.1016/j.jacc.2011.05.013

25. Wohlschlaeger J, Levkau B, Brockhoff G, et al. Hemodynamic support by left ventricular assist devices reduces cardiomyocyte DNA content in the failing human heart. Circulation. 2010;121(8):989–996. doi:10.1161/CIRCULATIONAHA.108.808071

26. Suzuki R, Li TS, Mikamo A, et al. The reduction of hemodynamic loading assists self-regeneration of the injured heart by increasing cell proliferation, inhibiting cell apoptosis, and inducing stem-cell recruitment. J Thorac Cardiovasc Surg. 2007;133(4):1051–1058. doi:10.1016/j.jtcvs.2006.12.026

27. A. Uygur RL. Mechanisms of Cardiac Regeneration. Dev Cell. 2017;36(4):362–374. doi:10.1016/j.devcel.2016.01.018.

28. Heallen T, Zhang M, Wang J, et al. Hippo pathway inhibits Wnt signaling to restrain cardiomyocyte proliferation and heart size. Science (80-). 2012;332(6028):458–461. doi:10.1126/science.1199010.Hippo

29. Xin M, Kim Y, Sutherland LB, et al. Hippo pathway effector Yap promotes cardiac regeneration. Proc Natl Acad Sci U S A. 2013;110(34):13839–13844. doi:10.1073/pnas.1313192110

30. Klotz S, Danser a. HJ, Foronjy RF, et al. The Impact of Angiotensin-Converting Enzyme Inhibitor Therapy on the Extracellular Collagen Matrix During Left Ventricular Assist Device Support in Patients With End-Stage Heart Failure. J Am Coll Cardiol. 2007;49(11):1166–1174. doi:10.1016/j.jacc.2006.10.071

31. Drakos SG, Drakos, S. G., Kfoury, A. G., Hammond, E. H., Reid, B. B., Revelo, M. P., Rasmusson, B. Y., … Li, D. Y. (2010). Impact of mechanical unloading on microvasculature and associated central remodeling features of the failing human heart. Journal of the Americ AG, Hammond EH, et al. Impact of mechanical unloading on microvasculature and associated central remodeling features of the failing human heart. J Am Coll Cardiol. 2010;56(5):382-391. doi:10.1016/j.jacc.2010.04.019

32. Xydas S, Rosen RS, Pinney S, et al. Reduced Myocardial Blood Flow During Left Ventricular Assist Device Support: A Possible Cause of Premature Bypass Graft Closure. J Hear Lung Transplant. 2005;24(11):1976–1979. doi:10.1016/j.healun.2005.03.003

33. Tansley P, Yacoub M, Rimoldi O, et al. Effect of left ventricular assist device combination therapy on myocardial blood flow in patients with end-stage dilated cardiomyopathy. J Heart Lung Transplant. 2004;23(11):1283–1289. doi:10.1016/j.healun.2003.09.005

34. Zhou P, Pu WT. Recounting cardiac cellular composition. Circ Res. 2016;118(3):368–370. doi:10.1161/CIRCRESAHA.116.308139

35. Weber KT, Sun Y, Bhattacharya SK, Ahokas R a, Gerling IC. Myofibroblast-mediated mechanisms of pathological remodelling of the heart. Nat Rev Cardiol. 2013;10(1):15–26. doi:10.1038/nrcardio.2012.158

36. Abbott CP, Lindsey ES, Creech O, Dewitt CW. a Technique for Heart Transplantation in the Rat. Arch Surg. 1964;89:645–652. doi:10.1001/archsurg.1964.01320040061009

37. Ono K, Lindsey ES. Improved technique of heart transplantation in rats. Transplantation. 1969;8(3):294. doi:10.1097/00007890-196909000-00022

38. Tevaearai HT, Walton GB, Eckhart AD, Keys JR, Koch WJ. Heterotopic transplantation as a model to study functional recovery of unloaded failing hearts. J Thorac Cardiovasc Surg. 2002;124(6):1149–1156. doi:10.1067/mtc.2002.127315

39. Ibrahim M, Navaratnarajah M, Kukadia P, et al. Heterotopic abdominal heart transplantation in rats for functional studies of ventricular unloading. J Surg Res. 2013;179(1):e31–e39. doi:10.1016/j.jss.2012.01.053

40. Fu X, Segiser A, Carrel TP, Tevaearai Stahel HT, Most H. Rat Heterotopic Heart Transplantation Model to Investigate Unloading-Induced Myocardial Remodeling. Front Cardiovasc Med. 2016;3(October). doi:10.3389/fcvm.2016.00034

41. Tanaka R, Tainaka M, Ota T, et al. Accurate determination of S-phase fraction in proliferative cells by dual fluorescence and peroxidase immunohistochemistry with 5- bromo-2’-deoxyuridine (BrdU) and Ki67 antibodies. J Histochem Cytochem. 2011;59(8):791–798. doi:10.1369/0022155411411090

42. Bologna-Molina, et al. Comparison of the value of PCNA and Ki-67 as markers of cell proliferation in ameloblastic tumors. Med Oral Patol Oral Cir Bucal. 2013;18(2). doi:10.4317/medoral.18573

43. Haubner BJ, Brice MA, Khadayate S, Metzler B, Aitman T, Penninger JM. Complete cardiac regeneration in a mouse model of myocardial infarction. Aging (Albany NY). 2012;4(12):966–977.

44. Sturzu AC, Rajarajan K, Passer D, et al. Fetal Mammalian Heart Generates a Robust Compensatory Response to Cell LossCLINICAL PERSPECTIVE. Circulation. 2015;132(2):109–121. doi:10.1161/CIRCULATIONAHA.114.011490

45. Naqvi N, Li M, Calvert JW, et al. A Proliferative Burst during Preadolescence Establishes the Final Cardiomyocyte Number. Cell. 2014;157(4):795–807. doi:10.1016/j.cell.2014.03.035

46. Xin M, Kim Y, Sutherland LB, et al. Hippo pathway effector Yap promotes cardiac regeneration. Proc Natl Acad Sci U S A. 2013;110(34):13839–13844. doi:10.1073/pnas.1313192110

47. Yeo MK, Kim SH, Kim JM, et al. Correlation of expression of phosphorylated and non-phosphorylated yes-associated protein with clinicopathological parameters in esophageal squamous cell carcinoma in a korean population. Anticancer Res. 2012;32(9):3835–3840.

48. Bassat E, Mutlak YE, Genzelinakh A, et al. The extracellular matrix protein agrin promotes heart regeneration in mice. Nature. 2017. doi:10.1038/nature22978

49. Schaefer A, Schneeberger Y, Schulz S, et al. Analysis of fibrosis in control or pressure overloaded rat hearts after mechanical unloading by heterotopic heart transplantation. Sci Rep. 2019;9(1):1–12. doi:10.1038/s41598-019-42263-1

50. Bonnet G, Zhao D, Wollmuth JR, et al. 100.59 High-Risk Percutaneous Coronary Intervention With or Without Prophylactic Mechanical Circulatory Support: Will Impella Show Superiority in the PROTECT IV Randomized Trial? JACC Cardiovasc Interv. 2024;17(4):S16–S17. doi:10.1016/j.jcin.2024.01.112

51. Goldstein DJ, Soltesz E. High-risk cardiac surgery : Time to explore a new paradigm. JTCVS Open. 2021;8(December):10–15. doi:10.1016/j.xjon.2021.10.001

